# ASS1 deficiency defines a therapeutic vulnerability in Philadelphia chromosome-positive acute lymphoblastic leukaemia

**DOI:** 10.64898/2026.05.21.724233

**Authors:** Michael J Austin, Shruti Patel, Ronas Kesmez, Foteini Kalampalika, Claudia Davies, Amit Sud, Aytug Kizilors, Nicholas Lea, Paulo Inocencio, Abigail Tappenden, Marianne Grantham, John Bomalaski, John Gribben, Miguel Ganuza Fernandez, Peter Szlosarek, Bela Patel

## Abstract

Amino acid deprivation with L-asparaginase is a cornerstone of treatment in acute lymphoblastic leukaemia (ALL), but clinical challenges limit its use in adults. Deficiency in the enzyme argininosuccinate synthase (ASS1-low) confers arginine auxotrophy, defining a dependence on extracellular arginine, and represents an analogue metabolic vulnerability that is targetable through arginine deprivation. We analysed transcriptomic data across >550 adult B-ALL cases to establish clinico-biologic characteristics of low *ASS1* expression identifying the Philadelphia chromosome-positive (Ph+) ALL subgroup as a stereotypical arginine auxotroph within molecularly diverse B-ALL. Functionally, arginine deprivation with pegargiminase induced robust apoptosis in both Ph+ ALL cell lines and primary samples in an ASS1-dependant manner and was highly effective as a monotherapy treatment in independent Ph+ ALL patient derived xenografts. Mechanistically, arginine deprivation induces endoplasmic reticulum stress mediated apoptosis which was orthogonal to tyrosine kinase inhibition (TKI) mediated outcomes. Using *in vitro* and *in vivo* models of non-genetically mediated TKI-resistance, we demonstrate pegargiminase and TKI combinations robustly eradicates TKI-resistant leukaemia. Thus, we establish ASS1 deficiency (ASS1-low) as a therapeutically actionable vulnerability in Ph+ ALL and a strategy to bypass TKI-resistance, supporting clinical evaluation of arginine deprivation as a novel adjunct to chemotherapy-free treatment.

## Introduction

Amino acid deprivation with asparaginase is a core component of multidrug treatment of Acute Lymphoblastic Leukaemia (ALL)^1^. Asparaginase targets the inability to endogenously synthesise asparagine due to suppression of the enzyme asparagine synthetase (ASNS) that renders cells reliant on extracellular asparagine and is a common feature of ALL blasts^2,3^. Several studies confirm the critical contribution of asparaginase to improved survival outcomes in paediatric and young adult ALL when incorporated into combination chemotherapy protocols^4–6^. However, its clinical utility is constrained by substantial toxicities including hepatotoxicity, pancreatitis and hypersensitivity reactions, with increased incidence and severity reported in adults^7^. Furthermore, variable intrinsic sensitivity to asparaginase is increasingly understood to be linked to molecular subtype^3,8^, highlighting the importance of cellular response determinants to therapeutic effectiveness. Together, these considerations underscore the importance of clinico-biologic heterogeneity as a focus for refinement of therapies.

An alternative amino acid deprivation strategy targeting the semi-essential amino acid arginine has emerged in recent years and has been tested in a range of cancers^9–14^. Arginine deprivation therapy is analogous to asparaginase wherein deficiency in the enzyme argininosuccinate synthase (ASS1) creates a dependence on extracellular arginine due to inability to synthesise arginine intracellularly, a state referred to as *arginine auxotrophy*. Both transcriptional and epigenetic gene suppression are mechanistically linked to ASS1 silencing^15–17^, which confers cancer cell fitness by increasing aspartate availability to fuel nucleotide synthesis for cellular proliferation and growth^18^. Clinically, low ASS1 expression is employed as a predictive biomarker of an arginine auxotrophic phenotype^19,20^.

Two arginine depleting drugs have progressed to clinical trials: pegylated arginine deiminase (ADI-PEG20, pegargiminase; Polaris Pharmaceuticals Inc, San Diego, USA), and pegylated recombinant human arginase (BCT-100; Bio-Cancer Treatment International, Hong Kong). Arginine deprivation agents are well tolerated even in older patient cohorts^9,13,14,19^, and in the phase III ATOMIC-MESO trial the addition of pegargiminase to frontline chemotherapy improved survival for patients with non-epithelioid mesothelioma^14^. However, responses across trials have varied, with greatest benefit found in tumours with the lowest expression of ASS1^19,20^ and when delivered as part of combination treatment^12,14^.

Susceptibility to arginine deprivation has previously been documented in a range of unselected ALL tumours^21,22^. However, an in-depth assessment of the potential for arginine deprivation to deliver targeted benefit across the full clinico-biological spectrum of ALL has not been performed.

Here we investigate predictive features for prospective identification of arginine auxotrophy in B-ALL and perform a systematic pre-clinical investigation of arginine deprivation therapy. Our findings support arginine auxotrophy as a class specific metabolic vulnerability of Philadelphia chromosome-positive (Ph+) ALL. Arginine deprivation treatment in this context yields anti-tumour effects orthogonally to standard-of-care tyrosine kinase inhibitor (TKI) treatment, thus providing a unique opportunity to expand current chemotherapy-free treatment paradigms in this setting.

## Methods

### Transcriptome analysis

RNA-sequencing HTSeq read counts from the study of Gu *et al*.^23^ were obtained from the St. Jude Cloud^24^ (https://www.stjude.cloud). RNA-sequencing BAM files from the study of Kim et al.^25^ were accessed via the European Genome-phenome Archive (dataset ID: EGAS00001007167). Gene expression microarray data from the study of Duy et al.^26^ were obtained from NCBI Gene Expression Omnibus (dataset ID: GSE23743). Developmental subclass of Ph+ ALL was assigned using the ALLCatchR algorithm^27^.

### Cell lines and culture conditions

Cell lines were obtained from DSMZ or ATCC. The p185^+^Arf^-/-^ cell line was kindly donated by the laboratories of Martine Roussel and Charles Scherr (St. Jude Children’s Research Hospital)^28–30^. Cells were maintained in culture for no more than 2 months.

### Primary cell culture

Primary human leukaemia cells were sourced from the HTA-licensed Barts Tissue Bank in compliance with approved local ethical protocols, as previously described^31^. Cells were cultured in a co-culture system using a GFP+ hTERT-immortalised human mesenchymal stem cell (MSC) feeder layer, given as a kind gift by Professor Dr D. Campana (St. Jude Children’s Hospital)^32^.

### Animal experiments

Experiments involving mice were performed with Queen Mary University of London veterinary oversight and complied with UK Home Office licensing requirements. C57BL/6 and NSG (NOD.Cg-Prkdc^scid^Il2rg^tm1Wjl^/SzJ) mice were purchased from Charles River Laboratories and housed in pathogen free conditions. Mixed sex mice were used for patient derived xenograft (PDX) experiments, while male mice were used for the p185^+^Arf^-/-^ dasatinib resistance model, based on previous engraftment and toxicity studies.

### *ABL1* kinase domain sequencing

*ABL1* kinase domain sequencing was performed as previously described^33^. Total RNA was extracted from mouse spleens and reverse transcribed to cDNA. The BCR::ABL1 transcript was then amplified using e1a2 specific primers. Library preparation was performed with the Illumina Nextera XT kit and paired-end sequencing was performed on an Illumina MiSeq platform. An in-house bioinformatics pipeline was utilised with a custom variant scoring tool for *ABL1* kinase domain variant identification.

Further details and additional methods can be found in supplementary methods.

## Results

### Reduced *ASS1* expression is characteristic of Ph+ ALL

Best responses to arginine deprivation have been reported in tumour subgroups with the lowest ASS1 expression^19,20^. However, meaningful biological differences are likely to exist outside a single selective cut-off^34^ and RNA based biomarkers using gene expression are challenging to implement clinically, prompting us to explore alternative predictors of arginine auxotrophy. We first analysed the distribution of *ASS1* expression within B-ALL using a publicly available RNA-sequencing dataset containing 501 individually profiled diagnostic adult ALL tumours^23^. *ASS1* expression displayed a positively skewed, multimodal distribution (Fig. 1a), suggestive of discrete levels of expression within adult B-ALL. We asked if these reflected subtype-specific *ASS1* expression levels and confirmed statistically significant differences between molecular subcategories of ALL (p =8.677 x10^-14^, Kruskal–Wallis test; Fig. 1b, S1). Among the most common B-ALL subtypes (>20 cases represented), Ph+ and Ph-like ALL exhibited the lowest median *ASS1* expression (Fig. 1b) and were therefore nominated as putatively arginine auxotrophic. We focused subsequent analyses on Ph+ ALL given the challenge of conventional asparaginase-based amino acid depletion strategies in this group^7^ and the relative uniformity in genetic background for evaluating subclass specific treatment effects.

**Figure 1:**
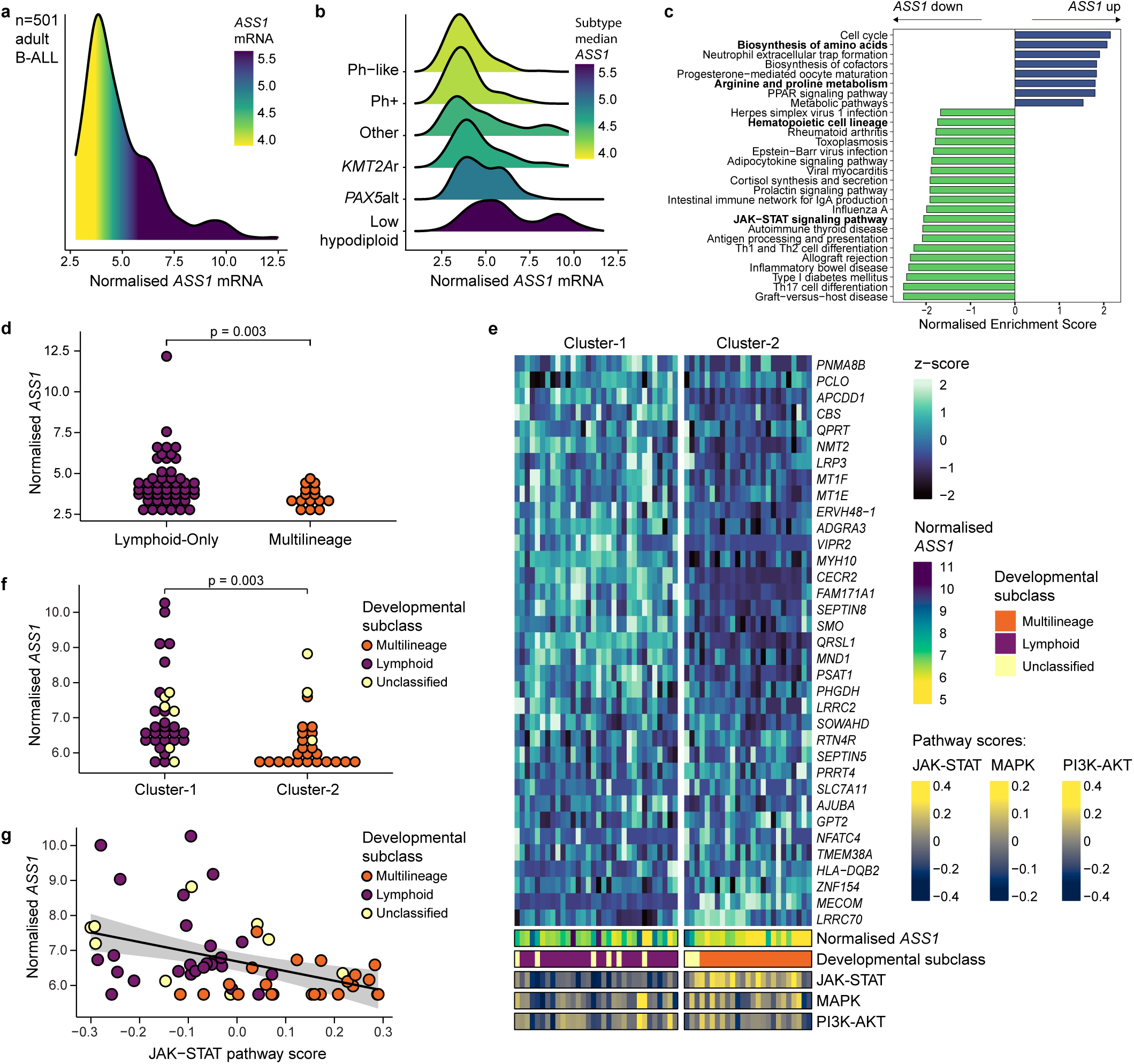
*ASS1* is suppressed in Ph+ ALL as part of coordinated transcriptional changes **(a)** Probability density function of normalised (*vst* transformed raw counts) *ASS1* expression across 501 adult patients (age >25 years at diagnosis) from the Gu *et al.* cohort^23^. **(b)** Subgroup-specific probability density functions of normalised *ASS1* expression from panel (a). Subgroups comprising >20 cases are shown; all subgroups are presented in Supplemental Fig. S1. Differences in distribution of *ASS1* ranks across subgroups were assessed using the Kruskal–Wallis test, p <0.0001. **(c)** Normalised enrichment scores for KEGG pathways with FDR <0.05 among *ASS1* co-expression signature genes in Ph+ sub-cohort from Gu *et al.*^23^. **(d)** Distribution of normalised *ASS1* expression between Lymphoid-Only and Multilineage developmental subclass cases in the Gu *et al.* cohort^23^. p-value from Welch’s t-test. **(e)** Consensus hierarchical clustering of Kim *et al.* cohort^25^ cases based on expression of *ASS1* co-expression signature genes with absolute Spearman’s ρ >0.4 (see Supplemental table ST1). **(f)** Distribution of normalised *ASS1* expression between Cluster-1 and Cluster-2 from Fig. 1e, coloured by developmental subclass. p-value from Welch’s t-test. **(g)** Correlation between normalised *ASS1* expression and JAK-STAT pathway score, Pearson’s *r* =0.71, *p* =4.28 x10^-10^.

Given ASS1 deficiency in other reported arginine auxotrophic tumours occurs within a wider coordinated metabolic, transcriptional and cellular programme^35–37^, we explored the global context of *ASS1* expression in Ph+ ALL at the transcriptomic level. We first derived a Ph+ ALL specific *ASS1* co-expression signature by performing pair-wise gene correlations with *ASS1* (Supplemental table ST1) and used this to perform Gene Set Enrichment Analysis (GSEA) to identify key biological features linked with putative arginine auxotrophy. We noted prominent enrichment of multiple cellular metabolic pathways as the most recurrent biological processes enriched amongst genes positively correlated with *ASS1*, including “Biosynthesis of amino acids” (NES =2.077, FDR =0.001) and “Arginine and proline metabolism” (NES =1.808, FDR =0.025) (Fig. 1c). Conversely, pathways enriched amongst genes negatively correlated with *ASS1* included “JAK–STAT signalling pathway” (NES =−2.051, FDR =0.003), known to operate downstream of BCR::ABL1^38^, and “Haematopoietic cell lineage” (NES =-1.734, FDR =0.022), which has an established association with fundamental Ph+ ALL biology^39^. Analogous *ASS1* co-expression signatures created in Philadelphia-negative counterparts did not yield similar pathway associations (Supplemental tables ST2-4, Fig. S2), suggesting that in Ph+ ALL a distinct transcriptional phenotype accompanies low *ASS1* expression and predicted arginine auxotrophy.

Given the negative correlation of the “Haematopoietic cell lineage” program (Fig. 1c) with *ASS1*, we compared expression between the Multilineage and Lymphoid-Only developmental subclasses of Ph+ ALL^39^ after *in silico* assignment of developmental state to individual cases^27,40^. Multilineage cases exhibited significantly lower mean *ASS1* expression than Lymphoid-Only cases (p =2.66 x10^-3^, Welch’s t-test; Fig. 1d) and a narrower distribution. Of note, *ASS1* expression amongst Lymphoid-Only cases was positively skewed, with most cases also expressing low-range *ASS1*. Thus, *ASS1*-low arginine auxotrophy may be a cross-cutting feature of Ph+ developmental subclasses, but is consistently enriched amongst Multilineage cases.

To validate these initial observations, we performed consensus hierarchical clustering using a subset of genes from the *ASS1* co-expression signature with absolute correlation coefficients of >0.4 in an independent dataset of 57 Ph+ ALL RNA-sequencing samples^25^. Two stable clusters were identified (Fig. S3) with distinct *ASS1* expression and distributions of developmental subclass (Fig. 1e). Cluster-1 had significantly lower mean A*SS1* expression than Cluster-2 (p <0.001, Welch’s t-test; Fig. 1f) and, in line with *ASS1* expression patterns documented previously (Fig. 1d), comprised exclusively Multilineage cases, while Cluster-2 comprised exclusively Lymphoid-Only cases (*p* =8.40 × 10⁻¹³, Fisher’s exact test; Fig. 1e).

Next, given the negative correlation of “JAK–STAT signalling pathway” with ASS1 expression (Fig. 1c), we compared *ASS1* expression with calculated JAK-STAT activity in the independent dataset to further define its relevance. This revealed a significant inverse correlation between JAK-STAT activity and *ASS1* expression (Pearson’s *r* =−0.43, p =9.29 × 10⁻⁴; Fig. 1g), indicating a likely biological intersection between low *ASS1* linked arginine auxotrophy and cellular signalling (Fig. 1c). We further compared JAK-STAT activity scores between clusters, noting higher activity in the Multilineage enriched Cluster-1 compared to Lymphoid-Only enriched Cluster-2 (p <0.001, Welch’s t-test, Fig. S4) consistent with a previously reported association^25^. Importantly, additional BCR::ABL1 related pathway activity scores did not correlate with *ASS1* expression (MAPK: Pearson’s *r* =−0.086, p =0.524; PI3K–AKT: Pearson’s *r* =-0.110, p =0.410; Fig. S5-6), in keeping with specific relevance of JAK-STAT signalling versus the broader BCR::ABL1 oncogenic programme to predicted arginine auxotrophy.

Finally, given the reportedly frequent occurrence of gene copy number variations (CNVs) in the flanking regions of the BCR::ABL1 translocation^41^, which includes the *ASS1* locus at 9q34.11, we utilised paired whole genome sequencing data^25^ to ask whether *ASS1* expression could be a function of *ASS1* copy number. We found that CNVs at the *ASS1* locus were distributed uniformly between developmental subclasses (p >0.99, Fisher exact test; Fig. S7), and there were no significant differences in *ASS1* expression between CNV states (p =0.577, Welch ANOVA; Fig. S8). We also compared pre-defined leukaemia associated gene CNVs from the same dataset^25^ with both cluster assignment and in line with their known co-linearity with developmental subclass^25,39^, we observed strong alignment of individual CNVs and the two identified clusters (Fig. S9). Given this co-linearity, we tested differences between *ASS1* expression and individual gene CNVs within each cluster and found no significant differences (Supplemental table ST5). These data suggest that variation in *ASS1* expression is not driven by *ASS1* copy number and is not associated with CNVs in common leukaemia associated genes independently of developmental subclass.

Altogether, these data identify Ph+ ALL as a B-ALL subtype with relatively low *ASS1* expression characterised by a putative arginine auxotrophy programme linked to developmental, metabolic and signalling states.

### ASS1 deficiency functionally sensitises Ph+ ALL to metabolic targeting

To formally assess whether ASS1 suppression in Ph+ ALL is a functional predictor of arginine auxotrophy, we evaluated sensitivity to arginine deprivation using the drug pegargiminase across a panel of Ph+ ALL cell lines spanning a range of baseline ASS1 expression (Fig. 2a). Pegargiminase induced a robust, dose-dependent decrease in leukaemia cell survival (Fig. 2b) together with substantial induction of apoptotic cell death (Fig. 2c-d) in ASS1-low cell lines (p185^+^*Arf*^−/−^ and TOM-1), whereas the ASS1-high cell line SUP-B15 was relatively resistant.

**Figure 2.**
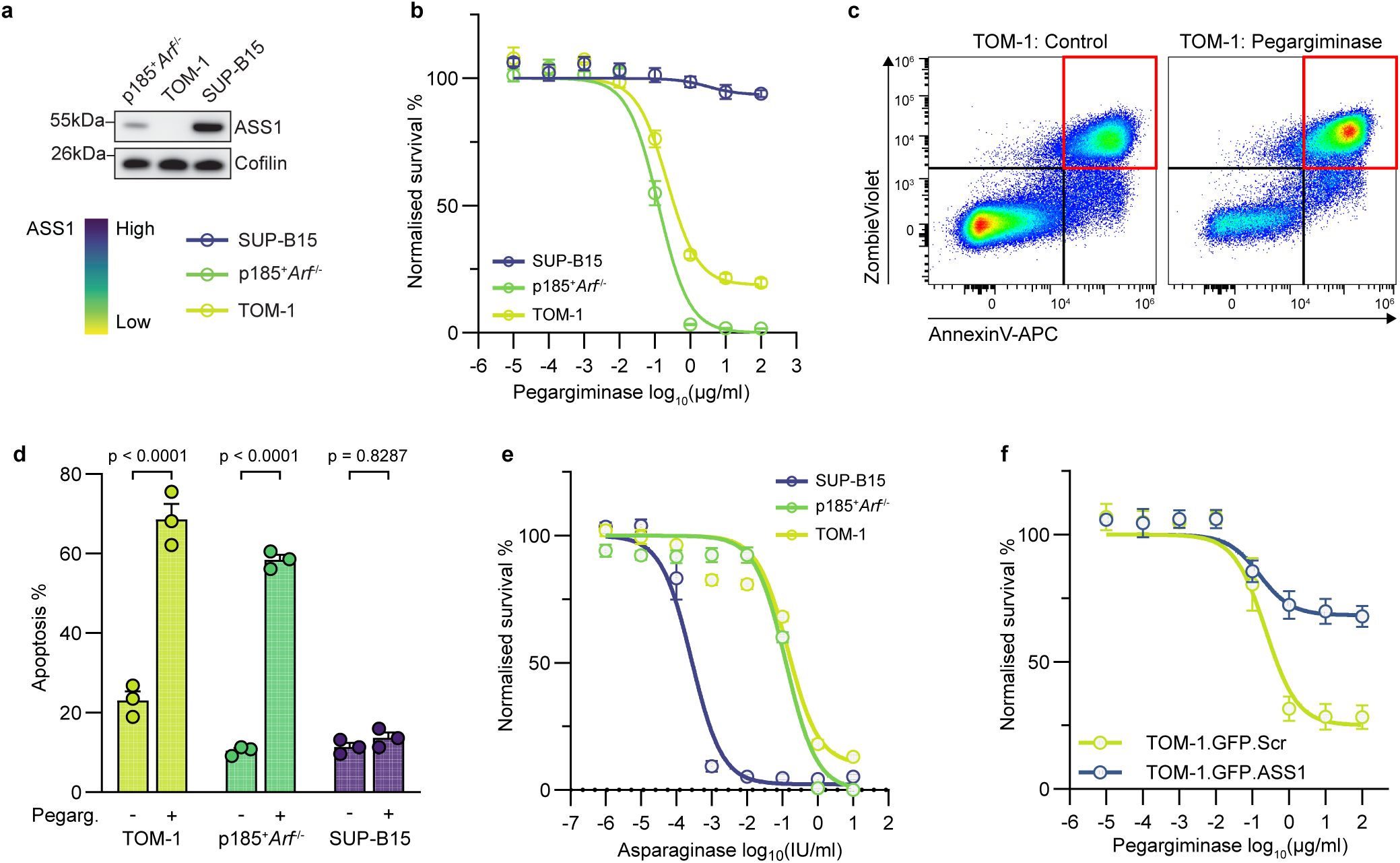
ASS1 deficiency defines a targetable metabolic vulnerability in Ph+ ALL. **(a)** Immunoblot analysis of baseline ASS1 protein expression in the indicated Ph+ ALL cell lines. Colour bar reflects qualitative ASS1 expression levels used in subsequent panels. **(b)** Dose–response curves for pegargiminase across Ph+ ALL cell lines, measured by ATP-based (CellTiter-Glo assay) estimation of live cell count and normalised to untreated controls. Line colour indicates baseline ASS1 expression as defined in panel (a). Points represent mean ± S.E.M. from n =3 independent experiments. **(c)** Representative flow cytometry density plots demonstrating pegargiminase-induced apoptosis in TOM-1 cells, with apoptotic cells highlighted in the Annexin-V+/Zombie-Violet+ (red) quadrant. **(d)** Quantification of apoptosis induction across Ph+ ALL cell lines following pegargiminase treatment, assessed by flow cytometry as in panel (e). Bar colour denotes baseline ASS1 expression as defined in panel (a). Points represent mean values from each independent experiment, error bars represent S.E.M. **(e)** Dose–response curves for asparaginase across Ph+ ALL cell lines, measured by ATP-based (CellTiter-Glo assay) estimation of live cell count and normalised to untreated controls. Line colour indicates baseline ASS1 expression as defined in panel (a). Points represent mean ± S.E.M. from n =3 independent experiments. **(f)** Dose–response curves for pegargiminase comparing TOM-1 cell line transduced with ASS1 cDNA or scramble control, measured by ATP-based (CellTiter-Glo assay) estimation of live cell count and normalised to untreated controls. Points represent mean ± S.E.M. from n =3 independent experiments.

To exclude the possibility of non-specific metabolic stress effects, we compared responses to asparaginase across the same cell lines. In line with pegargiminase targeting the specific dependency on extrinsic arginine, alternative amino acid withdrawal had relatively less impact in ASS1-low cell lines (Fig. 2e). To corroborate this principle, ASS1-low TOM-1 cells were virally transduced to ectopically express ASS1 (Fig. S10), which reduced susceptibility to pegargiminase (Fig. 2f) without altering response to asparaginase (Fig. S11), confirming ASS1-dependent anti-tumour responses to pegargiminase treatment.

To relate the functional relationship between variable ASS1 expression to earlier findings of JAK-STAT signalling, we assayed activation of STAT5, AKT and MAPK, representing canonical cascades downstream of BCR::ABL1. No apparent correlation was observed between baseline signalling pathway activity and ASS1 expression by immunoblotting (Fig. S12). Further assessments using pharmacological inhibition to modulate individual signalling cascades did not consistently modulate ASS1 expression, apart from in the ASS1-low p185^+^*Arf*^-/-^ cell line (Fig. S13), where MEK inhibition resulted in a robust reinduction of ASS1 expression, consistent with prior reports implicating RAS/MEK signalling in transcriptional suppression of ASS1 in other cancers^16^. Thus, ASS1 suppression in Ph+ ALL may involve context-dependent, BCR::ABL1 dependent and independent mechanisms.

Altogether, these results are consistent with ASS1-deficiency delineating therapeutically exploitable arginine auxotrophy, and provide confirmation of the utility of ASS1 expression for predicting pre-clinical efficacy of pharmacological arginine deprivation.

### Primary *ASS1*-low Ph+ ALL is susceptible to arginine deprivation *in vivo*

We next tested susceptibility to pegargiminase in a panel of five independent primary Ph+ ALL tumours (Supplemental table ST6) as a function of baseline *ASS1* expression using a stromal co-culture model system that supports primary leukaemia blast survival *ex vivo*^42^.

Pegargiminase reduced *ex vivo* leukaemic blast survival in four of five cases (“Responders”, Fig.3a), whereas resistance was observed in a singular case (ALL-04, Fig. 3a) which displayed distinctly higher baseline *ASS1* expression relative to the “responder” group.

**Figure 3:**
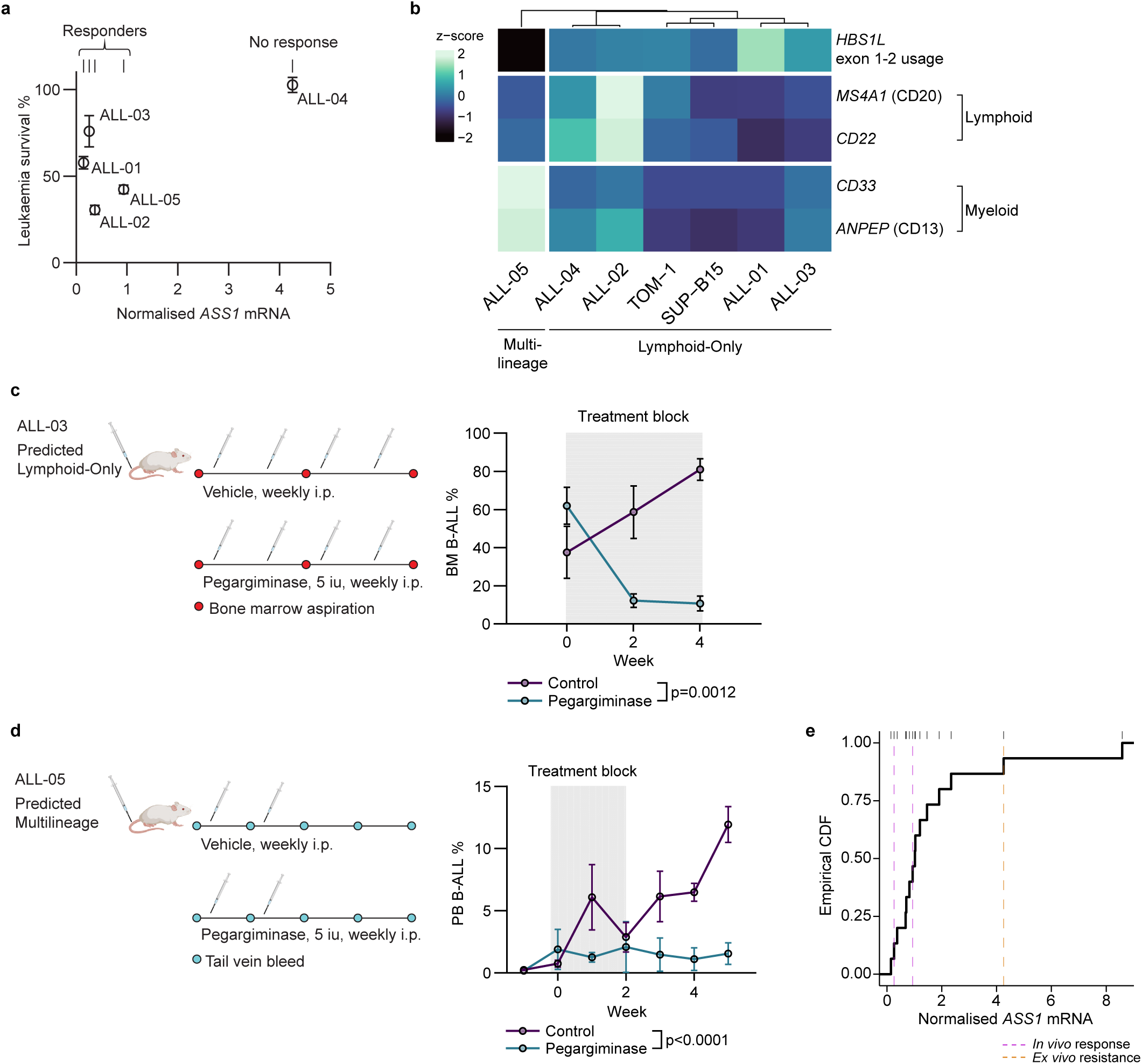
Pegargiminase is active in *ASS1*-low primary Ph+ ALL *in vivo*. **(a)** *Ex vivo* survival of primary Ph+ ALL leukaemic blasts following 48-hour pegargiminase exposure in hTERT-MSC co-culture, according to baseline *ASS1* mRNA expression. Viability was assessed by Annexin-V and viability dye co-staining after gating out GFP+ hTERT-MSC cells. Points represent mean ± S.E.M. from N =3 technical replicates for each primary sample. **(b)** Hierarchical clustering of Ph+ ALL primary and human cell line samples according to *HBS1L* exon 1-2 usage, expression of lymphoid genes *MS4A1* (CD20) and *CD22*, and expression of myeloid genes *ANPEP* (CD13) and *CD33*. Gene expression measured by RT-qPCR, with z-scores calculated per gene, from N =3 independent replicates per primary sample. **(c)** Treatment schema for the ALL-03 xenograft model and longitudinal assessment of bone marrow leukaemia burden by serial intra-tibial aspiration. Points represent mean ± S.E.M. from N =4 individual mice per arm. Statistical significance was evaluated using two-way ANOVA with treatment and time as factors. **(d)** Treatment schema for the ALL-05 xenograft model and longitudinal assessment of peripheral blood leukaemia burden by serial tail-vein sampling. Points represent mean ± S.E.M. from N =4 individual mice in pegargiminase group and N =3 mice in control group. Statistical significance was evaluated using two-way ANOVA with treatment and time as factors. **(e)** Empirical cumulative distribution function (CDF) of *ASS1* expression from N =10 diagnostic Ph+ ALL samples, including those used in panel (a). Dashed lines represent ASS1 expression levels of those samples tested *ex vivo* and/or *in vivo*, coloured by response type.

To determine if functional responses to pegargiminase are restricted to specific developmental subclasses, as potentially suggested by previously determined differences in *ASS1* expression between subclasses (Fig. 1d-e), we applied proxies of *HBS1L* exon 1-2 usage, expression of lymphoid-associated genes *MS4A1* (CD20) and *CD22*, and myeloid-associated genes *ANPEP* (CD13) and *CD33* ^25,39^ to assign a developmental subclass to each tested case (Fig. 3b). “Responders” were assigned to both Multilineage (ALL-05) and Lymphoid-Only (ALL-01, −02 and −03) subclasses, while the singular non-responding case ALL-04 was assigned to the Lymphoid-Only subclass. These results support low ASS1 expression as a functional predictor of susceptibility to arginine deprivation independent of developmental subclass.

To determine if *in vitro* pre-clinical responses to pegargiminase translate to an *in vivo* context, we generated PDXs by transplanting ALL-03 and ALL-05 into immunodeficient mice and then treated with 4 weekly doses or a truncated course of 2 weekly doses of pegargiminase, respectively. We observed a significant sustained monotherapy treatment effect irrespective of treatment schedule with statistically significantly lower leukemic burden in pegargiminase treated mice compared with vehicle treated controls (Fig. 3c-d).

To map these pegargiminase responses to a wider context, we modelled the distribution of expression across all *in vivo* (ALL-03 and ALL-05) and *ex vivo* (ALL-01, ALL-02 and ALL-04) assessed cases along with a second independent cohort of 10 diagnostic Ph+ ALL samples (Fig. 3e, Supplemental table ST6). Consistent with the RNA-sequencing assessment (Fig. 1b), *ASS1* expression demonstrated a strong positive skew, with the majority of cases falling within a narrow distribution containing the expression values of pegargiminase susceptible PDX cases (Fig. 3e). This therefore suggests that the majority of Ph+ ALL cases express *ASS1* in a range that would confer functional arginine auxotrophy.

### Arginine deprivation elicits antitumour effects independent of TKI

TKIs form the backbone of treatment of Ph+ ALL, including contemporary chemotherapy-free regimens incorporating immunotherapy into the frontline setting^43,44^.

We therefore evaluated pegargiminase integration with dasatinib, a contemporary second-generation TKI, across Ph+ ALL cell line models to establish potential to complement TKI response (Fig. 4a). In all models, the addition of pegargiminase to dasatinib reduced leukaemia cell survival compared with dasatinib alone (Fig. 4b). Of note, this effect was observed across all levels of ASS1 expression including the ASS1-high expressing SUP-B15 cell line where pegargiminase monotherapy was relatively ineffective (Fig. 2b). Similar additive effects were observed when combining pegargiminase to the first-generation TKI imatinib in the TOM-1 and p185^+^*Arf*^-/-^ cell lines, although the minor additive effect of pegargiminase in the ASS1-high SUP-B15 cell line was not statistically significant (Fig. S14).

**Figure 4:**
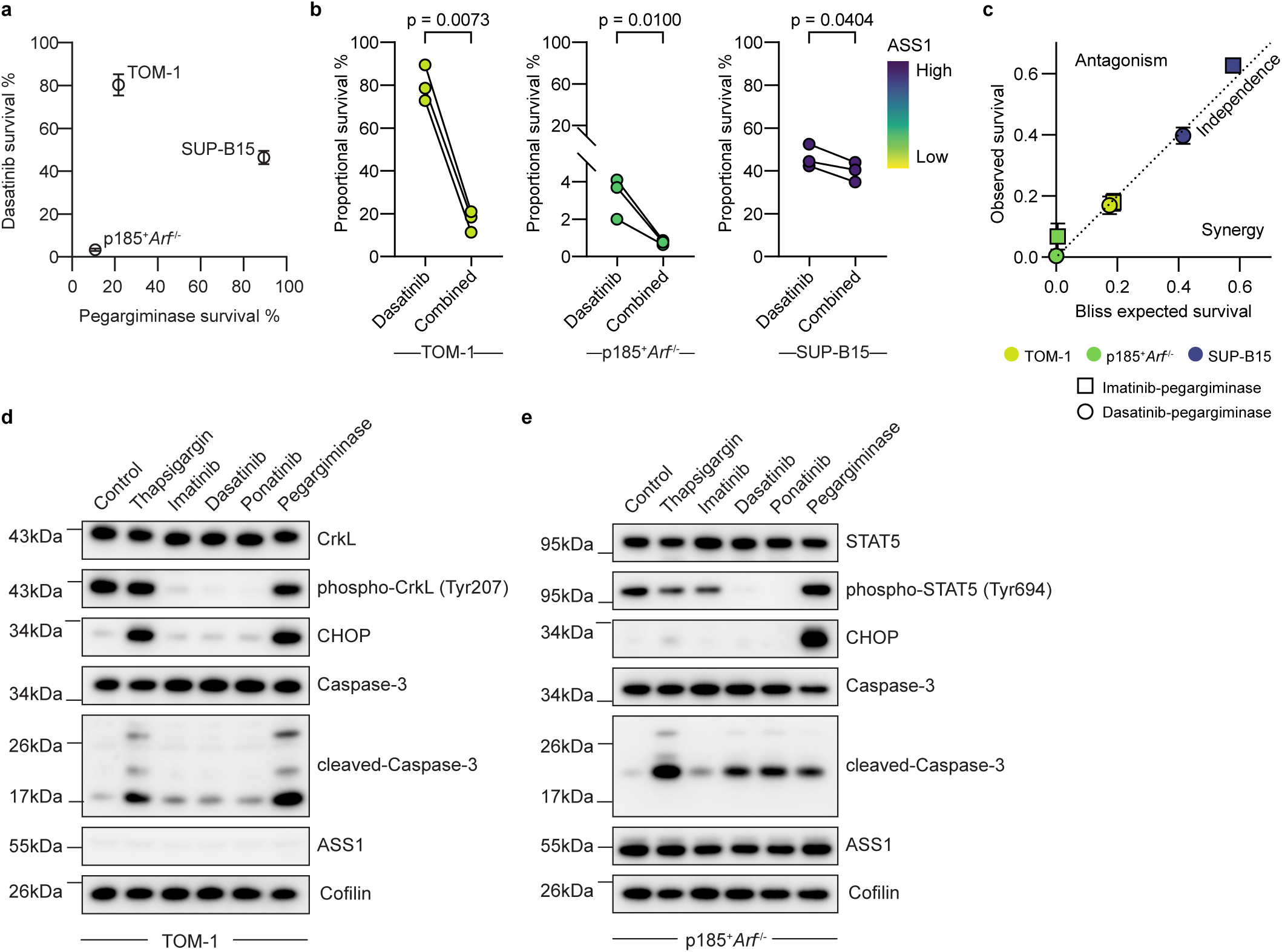
Pegargiminase exerts orthogonal effect to tyrosine kinase inhibition. **(a)** Relationship between survival following dasatinib monotherapy and survival following pegargiminase monotherapy across Ph+ ALL cell lines, assessed by Annexin-V and viability dye co-staining. Points and error bars represent mean and S.E.M. from N =3 independent experiments, respectively. **(b)** Comparison of survival following combined dasatinib and pegargiminase treatment versus dasatinib monotherapy across Ph+ ALL cell line panel. Point colour indicates baseline ASS1 expression as defined in Fig. 2a. Each point-pair represents paired mean survival values from independently performed experiments. Differences were assessed using paired ratio t-tests. **(c)** Bliss independence analysis comparing expected versus observed survival for combined dasatinib-pegargiminase and imatinib-pegargiminase treatments. Expected survival was calculated as the product of fractional monotherapy effects under an assumption of independent action. **(d)** Immunoblot analysis in TOM-1 cell line of CrkL dephosphorylation, CHOP induction, caspase-3 cleavage and ASS1 stability following 24-hour exposure to imatinib (4000 nM), dasatinib (100 nM), ponatinib (40 nM), pegargiminase (1000 ng/mL), or thapsigargin (250 nM). **(e)** Immunoblot analysis in p185^+^*Arf*^-/-^ cell line of STAT5 dephosphorylation, CHOP induction, caspase-3 cleavage and ASS1 stability following 24-hour exposure to imatinib (1000 nM), dasatinib (5 nM), ponatinib (20 nM), pegargiminase (1000 ng/mL), or thapsigargin (20 nM).

Using the Bliss independence model to understand the basis of the combination effect we compared observed survival with calculated expected survival for combination treatments in each cell line. Across all models, observed survival closely matched that expected, suggestive of independent, additive effects (Fig. 4c). To explore this at a mechanistic level, we compared global transcriptomic changes induced by pegargiminase using RNA-sequencing of the pegargiminase-sensitive, ASS1-low TOM-1 cell line with publicly available transcriptomic data from imatinib-treated Ph+ ALL cell lines^26^ and found minimal overlap in gene expression perturbation between drugs (Fig. S15), supportive of mechanistically orthogonal drug lethality.

To further investigate this at the molecular level, we examined apoptosis cascades amongst differentially expressed genes from the ASS1-low TOM-1 cell line RNA-sequencing and found multiple pro-apoptotic genes induced by pegargiminase involving the endoplasmic reticulum (ER) stress response, including CHOP (*DDIT3*), DR5 (*TNFRSF10B*), GADD34 (*PPP1R15A*) and TRIB3 (Fig. S16), consistent with known cellular responses to amino acid deprivation^45,46^. Given the central role of CHOP in instigating ER-stress mediated apoptosis^45^, we directly assayed CHOP induction following pegargiminase and TKI treatment. Consistent with orthogonal mechanisms, TKI induced cleavage of caspase-3 with demonstratable inhibition of BCR::ABL1 signalling and without CHOP induction, whereas pegargiminase induced cleavage of caspase-3 without inhibition of BCR::ABL1 signalling but with potent induction of CHOP, matching the pattern of the ER-stressor positive control thapsigargin (Fig. 4d-e). Importantly, we observed no apparent modulation of ASS1 expression by TKI treatment, further supporting non-overlapping, pharmacologically orthogonal pathway activity between pegargiminase and TKI.

### Metabolic targeting of ASS1 deficiency bypasses non-genetic resistance to TKI therapy

Later generation TKIs, including dasatinib and ponatinib, are known to reduce the incidence of *ABL1* kinase domain mutation mediated, genetic TKI-resistance in Ph+ ALL^47^. Nevertheless, a substantial proportion of patients will fail to achieve deep molecular remissions at the end of contemporary chemotherapy-free induction therapy despite documented low rates of *ABL1* kinase domain mutation^43,44,47^. This suggests suboptimal disease response may derive from non-genetic TKI-resistance factors. Thus, targeting these BCR::ABL1 independent resistance mechanisms may further increase and deepen remission responses.

Given our demonstration that pegargiminase acts orthogonally to TKI (Fig. 4d-e, S15), we hypothesised that arginine deprivation using pegargiminase may be an effective strategy against non-BCR::ABL1 mediated TKI-resistance.

To formally test this, we employed the p185⁺*Arf*^⁻/⁻^ cell line, which acquires functional resistance to TKI in the presence of interleukin-7 (IL-7)^48^ via sustained STAT5 signalling (Fig. 5a), representing a well-established non-genetic resistance model that mirrors clinically reported resistance modes^49^. We observed increases in residual leukaemia cell survival following either dasatinib or ponatinib monotherapy in the presence of IL-7 (Fig. 5b), confirming model validity. In contrast, combining pegargiminase to TKI significantly reduced IL-7 driven TKI-resistant cells, consistent with the notion that combination treatments incorporating pegargiminase impact non-BCR::ABL1 driven TKI-resistance to augment anti-tumour response. (Fig. 5b).

**Figure 5.**
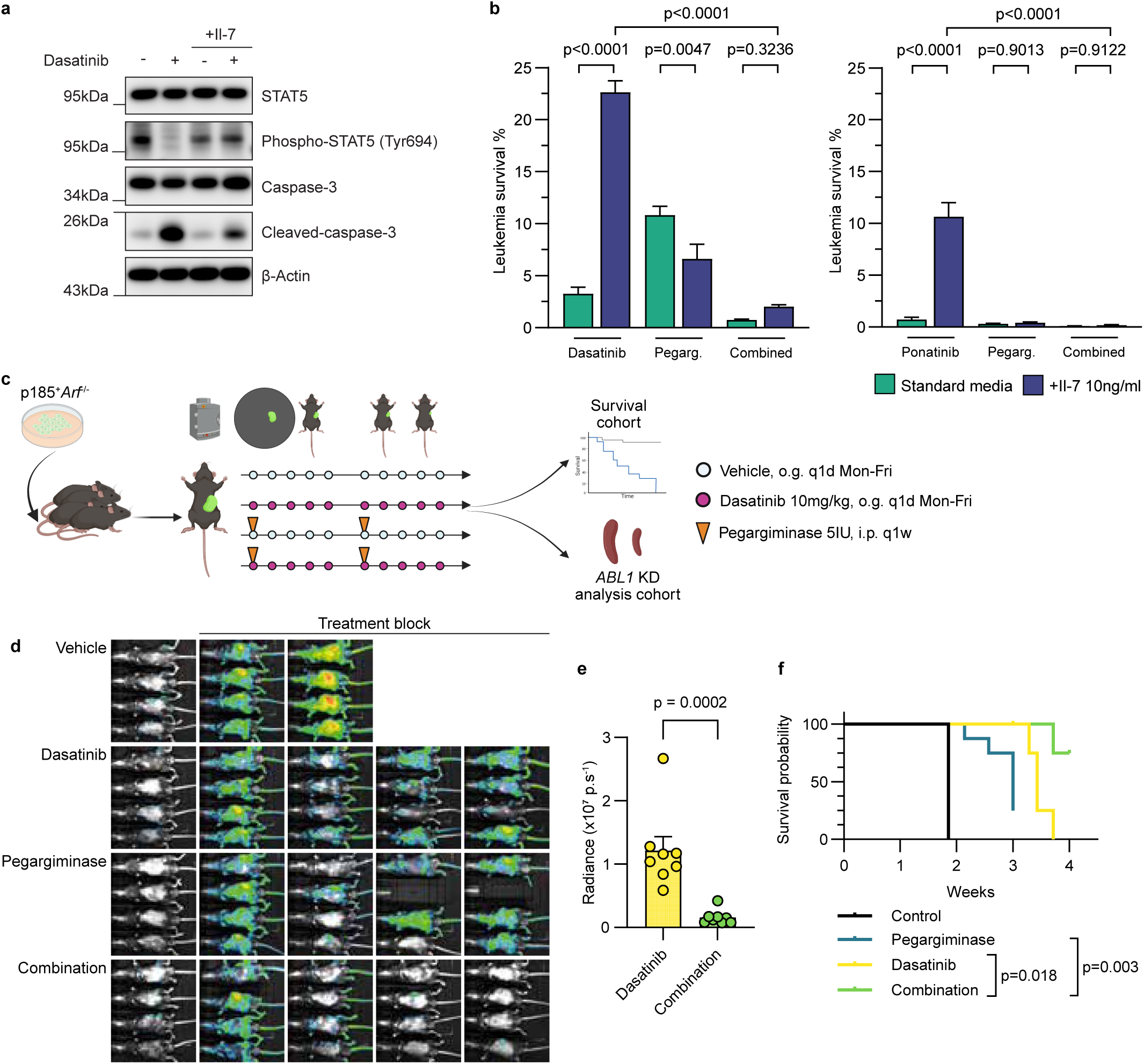
Pegargiminase overcomes non-genetic TKI-resistance and deepens therapeutic response. **(a)** Immunoblot analysis of STAT5 dephosphorylation and caspase-3 cleavage following 24-hour exposure to dasatinib (5 nM) with or without recombinant IL-7 (10 ng/mL). **(b)** Viability following 72-hour dasatinib treatment (5 nM, left panel), ponatinib treatment (2 nM, right panel), pegargiminase (1000 ng/mL) or combinations, with or without recombinant IL-7 (10 ng/mL). Viability was assessed by Annexin-V and viability dye co-staining. Bars and error bars represent mean and S.E.M. of N =3 independent experiments, respectively. Statistical significance was evaluated using two-way ANOVA with drug treatment and IL-7 exposure as factors. **(c)** Experimental schema for *in vivo* modelling of dasatinib resistance. **(d)** Representative merged bioluminescence and bright-field images from mice during treatment (n =8 per treatment arm; n =4 for vehicle controls). **(e)** End-of-treatment whole-body radiance quantification. Points represent individual mouse radiance values. Group differences were assessed by t-test. **(f)** Kaplan–Meier survival curves for mice in each treatment arm. Differences were assessed using the log-rank test with a significance threshold of p <0.025 to account for two comparisons.

We next modelled non-genetic resistance *in vivo*, using a dasatinib monotherapy schedule in a well-documented transgenic BCR::ABL1 model^30^ (Fig. 5c). Dasatinib monotherapy treated mice initially responded but then relapsed during the 2-week treatment block, simulating a clinically relevant dynamic of response followed by emergent TKI-resistance (Fig. 5d). Pegargiminase monotherapy initially delayed leukaemia progression (Fig. 5d), but 6/8 mice eventually displayed progressive leukaemia and were euthanised by the end of the 2-week treatment period. In contrast, the combination of pegargiminase and dasatinib resulted in markedly sustained disease control throughout the treatment period (Fig. 5d). Combination treatment recipients all survived to the end of treatment and were found to demonstrate a significant reduction in residual leukaemia by bioluminescence compared to dasatinib monotherapy treated mice (p =0.0002, t-test; Fig. 5e), as well as statistically significant prolongation in survival compared to either monotherapy arms (Fig. 5f).

Finally, we sequenced the *ABL1* kinase domain from residual blasts from n =4 randomly chosen mice from both dasatinib monotherapy and combination arms at the end of treatment, as well as n =3 control mice taken at the time of euthanasia. No detectable TKI-resistance mutations at a sensitivity level of 1% were found, validating these pre-clinical outcomes as bona fide demonstrations of treatment efficacy in a setting of phenotypic non-genetic TKI-resistance.

## Discussion

Personalised metabolic therapies may offer individualised, highly tailored approaches to improving outcomes at patient-level, a principle that has been shown in select contexts, including von Hippel Lindau disease related tumours^50^ and isocitrate dehydrogenase mutant acute myeloid leukaemia and gliomas^51,52^ but not yet substantively in ALL.

Using complementary analysis of large clinical cohort datasets and pre-clinical functional modelling, we identify ASS1-deficient arginine auxotrophy as a therapeutically exploitable vulnerability of Ph+ ALL. This constitutes the first notable demonstration of a class-specific metabolic vulnerability in ALL.

Recent clinical studies have demonstrated that chemotherapy-free treatment strategies combining later-generation tyrosine kinase inhibitors (TKIs) with immunotherapy achieve superior outcomes compared with traditional chemotherapy-based regimens in Ph+ ALL^43,44,53^. Nevertheless, a substantial proportion of patients fail to achieve deep molecular responses during induction, and relapse at extramedullary sites, including the central nervous system, is emerging as a key issue associated with such an approach^44,53^. Importantly, our data demonstrate that arginine deprivation can deepen responses to TKI therapy and maintain activity in models of non-genetic resistance *in viv*o, highlighting a potential approach to address residual disease in a chemotherapy-free context.

Long-term follow up data from current chemotherapy-free treated patients have shown that a complete molecular response after TKI induction is associated with 100% long term disease-free survival rates^54^, highlighting the importance of early response status. Thus, the addition of complementary, low-toxicity drugs early in treatment may increase deep molecular responses by the end of induction. Adjuncts reported to enhance TKI therapy, including arsenic trioxide, proteasome inhibitors and histone deacetylase inhibitors, have not progressed clinically^55–57^, and are not necessarily compatible with chemotherapy-free approaches. The demonstration that arginine deprivation using pegargiminase enhances TKI responses is therefore significant given the well-established clinical safety profile of pegargiminase across both solid and haematological malignancies, and in older patients^11,14^.

Consistent with observations in other tumour types, we show that sensitivity to pegargiminase in Ph+ ALL is driven by ASS1 deficiency. While ASS1 suppression has been reported to occur downstream of RAS/MEK signalling in non-small cell lung cancer^16^, we observed only partial concordance across Ph+ ALL models, indicating that ASS1 suppression may be context dependent and warrants further study, including epigenetic regulation. The strong association between lowest *ASS1* expression and the Multilineage developmental subclass also suggests that in this context *ASS1* suppression may reflect broader lineage or developmental programmes and may overlap mechanistically with the reported suppression of *ASS1* in Chronic Myeloid Leukaemia stem cells^58^. In our *in silico* analysis, we also uncovered a co-association with developmental subclass, increased JAK-STAT signalling and *ASS1* suppression, but were unable to systematically test this association given that mainly Lymphoid-Only cell line and primary models were employed.

Despite developmental subclass differences in *ASS1* expression, we demonstrated responses to pegargiminase in both Multilineage and Lymphoid-Only models and estimate that most Ph+ ALL cases express *ASS1* within a range conferring susceptibility to arginine deprivation, indicating that combining pegargiminase with contemporary Ph+ ALL treatment has the potential to benefit most patients. Identifying patients who would have the greatest benefit from the addition of arginine deprivation to treatment remains a challenge, and while Multilineage subclass related genomic features, such as monosomy 7 or deletion of *HBS1L*^39^, would have utility in identifying patients predicted to have consistently low *ASS1* expression, this would exclude patients with *ASS1*-low, Lymphoid-Only subclass disease. Therefore, the identification of prospective biomarkers that are robust and objective identifiers of arginine auxotrophy remains a priority.

A key focus of our study was the challenge of non-genetically mediated TKI-resistance, which is increasingly recognised as a contributor to treatment failure in Ph+ ALL^49^. While *ABL1* kinase domain mutation has long been recognised as the canonical mode of TKI-failure in Ph+ ALL, its clinical relevance has diminished with the upfront use of later-generation TKIs such as ponatinib^47^. Indeed, recent trials have reported very low rates of kinase domain mutations despite persistent measurable residual disease at the end of induction^44^. Our finding that arginine deprivation retains activity in models of non-genetic TKI-resistance and can significantly reduce residual disease burden when added to TKI treatment, therefore supports its potential role as a complementary strategy to deepen molecular responses.

Finally, chemotherapy-free regimens have exposed increased relapse rates in the CNS and other extramedullary sites. While we did not directly assess CNS activity in this study, pegargiminase has previously been shown to be active against tumours of the CNS^59^ and is currently in a phase 3 trial for patients with glioblastoma (NCT03970447), raising the possibility that arginine deprivation could contribute to CNS disease control when combined with TKI therapy, something that would require formal testing in future pre-clinical and clinical studies.

## Data availability

Primary RNA-sequencing data generated in this project have been deposited in NCBI’s Gene Expression Omnibus and are accessible through GEO Series accession number GSE330924 (https://www.ncbi.nlm.nih.gov/geo/query/acc.cgi?acc=GSE330924).

## Acknowledgements

The authors would like to thank staff from the Barts Cancer Institute tissue bank for sample collection and processing, and the staff from the Queen Mary University of London Biological Services Unit.

This work was supported by Blood Cancer UK (grant 15009; B Patel), The Greg Wolf Foundation (B Patel), Cancer Research UK (grant CRUK A21019; B Patel, co-applicant), City of London, CRUK (B Patel) and Gabrielle’s Angels Foundation (B Patel) and Barts Charity (grant G-00238; B Patel) and an NIHR clinical academic training grant (B Patel and M Austin).

## Authorship Contributions and Conflicts of Interest

**Michael J Austin**: conceptualisation, methodology, software, formal analysis, investigation, writing - original draft, writing - review & editing, visualisation. **Shruti Patel**: investigation, writing - review & editing. **Ronas Kesmez**: methodology, investigation, software, writing - review & editing. **Foteini Kalampalika**: methodology, writing - review & editing. **Claudia Davies**: methodology, writing - review & editing. **Amit Sud**: methodology, software, writing - review & editing. **Aytug Kizilors**: methodology, investigation, writing - review & editing. **Nicholas Lea**: methodology, investigation, writing - review & editing. **Paulo Inocencio**: methodology, investigation, writing - review & editing. **Abigail Tappenden**: resources, writing - review & editing. **Marianne Grantham**: resources, writing - review & editing. **John Bomalaski**: resources, writing - review & editing. **John Gribben**: resources, writing - review & editing. **Miguel Ganuza Fernandez**: resources, methodology, writing - review & editing. **Peter Szlosarek**: resources, methodology, writing - review & editing. **Bela Patel**: conceptualisation, methodology, writing - original draft, writing - review & editing, supervision, project administration, funding acquisition.

**Michael J Austin**: Polaris Pharmaceuticals, Research Funding. **Shruti Patel**: Polaris Pharmaceuticals, Research Funding. **John Bomalaski**: Polaris Pharmaceuticals, Current Employment. **Peter Szlosarek**: Polaris Pharmaceuticals, Research Funding. **Bela Patel**: Polaris Pharmaceuticals, Research Funding. All other authors declare no relevant conflicts of interest.

## Supplemental figure legends

**S1.** Subgroup-specific probability density functions of normalised *ASS1* expression from all subgroups with N >2 adult patients. Differences in distribution of *ASS1* ranks across subgroups were assessed using the Kruskal–Wallis test, *p* =8.68 x10^-14^.

**S2.** Normalised enrichment scores for KEGG pathways with FDR <0.05 among *ASS1* co-expression signature genes in *KMT2A*r, *PAX5*alt and Ph-like sub-cohorts from Gu *et al.*^23^.

**S3.** Consensus clustering outputs for hierarchical clustering in Fig. 1e. Left panel: Cumulative distribution functions (CDFs) of consensus index scores per iteration of cluster number, *k*. Right panel: Relative change in area under CDF curve per cluster *k* iteration.

**S4.** Distribution of JAK-STAT pathway scores between Cluster-1 and Cluster-2 from Fig. 1e, coloured by developmental subclass. p-value from Welch’s t-test.

**S5.** Correlation between normalised *ASS1* expression and MAPK pathway score, Pearson’s *r* 0.16, *p* =0.242.

**S6.** Correlation between normalised *ASS1* expression and PI3K-AKT pathway score, Pearson’s *r* 0.02, *p* =0.909.

**S7.** Area plot illustrating distribution of CNVs at *ASS1* locus between developmental subclasses.

**S8.** Distribution of normalised *ASS1* expression between *ASS1* copy number states. p-value from Welch’s ANOVA test.

**S9.** Oncoprint illustrating distribution of pre-annotated CNVs in leukaemia associated genes^25^ between clusters. Adjusted p-values from Fisher’s exact test of distribution of per-gene CNVs between clusters.

**S10.** Immunoblot analysis of ASS1 expression between TOM-1 cells transduced with ASS1 cDNA or scrambled construct.

**S11.** Dose–response curves for asparaginase comparing TOM-1 cell line transduced with ASS1 cDNA or scramble control, measured by ATP-based (CellTiter-Glo assay) estimation of residual live cell counts and normalised to untreated controls. Points represent mean ± S.E.M. from N =3 independent experiments.

**S12.** Immunoblots of baseline phosphorylation of signalling pathway elements from AKT, STAT5 and MAPK pathways across Ph+ ALL cell line panel.

**S13.** Immunoblot analysis of ASS1 expression upon inhibition of BCR::ABL1 with dasatinib and major downstream signalling components using ruxolitinib (JAK1/2), trametinib (MEK1/2) and LY294002 (PI3K) inhibitors.

**S14.** Comparison of survival measured by flow cytometry following combined imatinib and pegargiminase treatment versus imatinib monotherapy across a Ph+ ALL cell line panel. Point colour indicates baseline ASS1 expression as defined in Fig. 2a. Each point-pair represents paired mean survival results from independently performed experiments, each individually performed in technical triplicate. Differences were assessed using paired ratio t-tests.

**S15.** Overlap of differentially expressed genes between imatinib treated Ph+ ALL cell lines (N =4) from Duy et al.^26^, from microarray data analysis with nominal p-value threshold for significance of 0.05, and pegargiminase treated TOM-1 cell line from RNA-sequencing with FDR threshold for significance of 0.001.

**S16.** Volcano plot of differentially expressed genes between pegargiminase treated and control in TOM-1 cell line. Labelled genes are those associated with gene ontology pathway GO:0006915 “Apoptotic process” with log2 fold change in gene expression >2 and FDR <0.0001.

