## supplemental_methods for "ASS1 deficiency defines a therapeutic vulnerability in Philadelphia chromosome-positive acute lymphoblastic leukaemia"

### Transcriptome analysis

RNA-sequencing HTSeq read counts from the study of Gu *et al*.^1^ were obtained from the St. Jude Cloud^2^ (<https://www.stjude.cloud>) and batch corrected using the ComBat function from the R package sva^3^. RNA-seq BAM files from the study of Kim et al.^4^ were accessed through the European Genome-phenome Archive (dataset ID: EGAS00001007167). Gene-level counts were generated with Rsubread^5^ using Queen Mary University of London's (QMUL) Apocrita HPC facility, supported by QMUL Research-IT^6^. Gene expression microarray data from the study of Duy et al.^7^ were obtained from NCBI Gene Expression Omnibus (dataset ID: GSE23743).

Developmental subclass of Ph+ ALL was assigned using the ALLCatchR algorithm using raw RNA-sequencing counts^8^. Microarray and RNA-seq data were analysed with R packages limma^9^ and DESeq2^10^, respectively. For visualisation of RNA-seq data, the DESeq2 *vst* transformation was applied to raw counts.

The *ASS1* co-expression signature was generated following the method of Chan et al.^11^. Spearman gene-wise expression correlation coefficients were calculated using the psych package^12^ after filtering the data to include the 5000 top genes ranked by median absolute deviation of expression. Gene set enrichment analysis was performed using the clusterProfiler package^13^ with pathway definitions from the Kyoto Encyclopedia of Genes and Genomes (KEGG). Consensus hierarchical clustering was performed using the package ConsensusClusterPlus^14^. Data representations were generated with ComplexHeatmap^15^ and tidyverse packages^16^.

To generate primary RNA-sequencing data, total RNA was extracted from cell lines the QIAgen RNeasy kit following the manufacturer’s instructions. RNA concentration and integrity were assessed using the Agilent Tapestation system. Poly(A) tail capture based library preparation was then performed, followed by 150bp paired end sequencing with 20M reads/sample using the Illumina NovaSeq platform. Raw reads were aligned to the hg38 reference genome with STAR^17^, using the QMUL Apocrita HPC facility^6^, and then further processed as described above.

### Copy number analysis

Aligned BAM files were downloaded from the European Genome Phenome Archive using pyEGA3 (v5.2.0)^18^ and realigned to GRCh38 (with alt and HLA contigs^19^) using Bazam (v1.1.0)^20^. Resulting realigned BAM files were sorted with Samtools (v1.19.2)^21^ and duplicate reads marked and removed with Sambamba (v1.0.0)^22^.

Genome-wide copy-number alterations were called using ichorCNA (v0.3.1)^23^. 10 kb-bin read count files were generated with readCounter from the HMMcopy suite (v1.5.2)^24^. ichorCNA was run with default parameters, with the exception of estimateScPrevalence=FALSE and the inclusion of chromosome Y in the analysis. As ichorCNA purity estimates were inconsistent across samples, tumour purity was assigned from whichever of three sources best agreed with the ichorCNA logR plots: ichorCNA itself, PURPLE (v3.9.2)^25^, or blast percentage estimates from the original publication^4^. We classified 10 kb bins of logR values (extracted from .cna.seg files generated by ichorCNA) across genes of interest as homozygous deletions, heterozygous deletions, gains, or amplifications. Custom purity values could not be imputed into ichorCNA, so we adapted a purity and ploidy-adjusted logR equation from the CNVKit package^26^ to classify each bin, except for homozygous deletions, where a hardcoded logR<-2.0 cut-off was implemented. CNV concordance across IKZF1, CDKN2A, and HBS1L was calculated against the original paper^4^. We observed high concordance in deletion detection (≥89% concordance across all types of CNVs). For analysis of CNVs of leukaemia associated genes defined by the original authors^4^, pre-annotated CNV data were taken directly from the original publication.

### Cell lines and culture conditions

Cell lines were obtained from DSMZ or ATCC. The p185^+^Arf^-/-^ cell line was kindly donated by the laboratories of Martine Roussel and Charles Scherr (St. Jude Children’s Research Hospital)^27–29^. TOM-1 and SUP-B15 cell lines were cultured in RPMI media supplemented with 20% FBS and 1% penicillin-streptomycin in a standard humidified incubator. The p185^+^Arf^-/-^ cell line was cultured in RPMI media supplemented with 10% FBS, 1% penicillin-streptomycin and 1X β-mercaptoethanol (Gibco).

Cells were passaged by centrifugation and maintained in culture for no more than 2 months after first thawing. Cultures were routinely tested for mycoplasma contamination.

### Drug treatments

Drugs were added to cells growing in log-phase at concentrations indicated in relevant figures. Control conditions used matched DMSO concentration, where appropriate. Pegargiminase was obtained from Polaris Pharmaceuticals Inc, in collaboration with Professor Peter Szlosarek (QMUL). Imatinib was purchased from Scientific Laboratory Supplies. Dasatinib, E. Coli expressed asparaginase and thapsigargin were purchased from Merck. LY294002, ponatinib, trametinib and ruxolitinib were purchased from Cambridge Bioscience.

Responses to drug concentration titrations were assessed using the CellTiter-Glo assay (Promega), following the manufacturer’s instructions. Briefly, cells were seeded in opaque white 96-well plates then following drug treatments cells were lysed with the CellTiter-Glo reagent and luminescence was measured with an FLUOstar Omega plate reader (BMG Labtech). Luminescence values were normalised per row and then averaged over triplicate repeats per plate. Dose-response curves represent mean values from at least 3 independent plates, generated using non-linear regression in GraphPad Prism.

For fixed dose TKI experiments, concentrations for human cell lines were selected to reflect clinically relevant drug exposures based on published pharmacokinetic data^30–33^. Concentrations for the mouse cell line p185^+^Arf^-/-^ were based on titrations experiments.

### Flow cytometry

Flow cytometry analysis was performed as previously described^34^ using an Attune NXT flow cytometer (Thermo Fisher) and analysed with FlowJo software (version 10, Beckton Dickinson).

For *in vitro* drug treatment assessment, cells were stained with Zombie Violet viability dye (BioLegend) and AnnexinV-APC (BioLegend). Single-stained cells were used to compensate for spectral overlap and define gate thresholds. Debris and doublet cells were gated out using forward- and side-scatter channels. Apoptotic/dead and residual live cells were quantified as double-positive or double-negative populations, respectively.

For *in vivo* analysis of mouse blood or bone marrow, samples were washed and red blood cells were lysed using RBC Lysis Buffer (BioLegend). Cells were stained with Zombie Violet viability dye and a panel of antibodies against mouse-CD45-FITC (BioLegend), human-CD45-APC (BioLegend) and human-CD19-PE-Cy5.5 (BioLegend). Single stained compensation beads were used to correct spectral overlap and flow-minus-one controls were used to set gate thresholds. Leukaemia burden was quantified as the ratio of human-CD45+/human-CD19+ cells to total leukocytes, quantified as combined mouse-CD45+ and human CD45+ populations.

### ASS1 overexpression

### ASS1-overexpressing TOM-1 cells were generated using commercially available lentiviral particles encoding either a GFP-tagged ASS1 cDNA clone (OriGene, catalogue number: RC201130L2V) or GFP-tagged control cDNA sequence (OriGene, catalogue number: PS100071V). Parental TOM-1 cells were seeded in non-tissue culture treated plates after pre-impregnated with 12μg/ml retronectin solution for 24 hours. Cells growing in logarithmic phase were then transduced with lentiviral particles at a Multiplicity of Infection (MOI) of at least 5. After 3 days, cells were passaged with fresh media. Stable transgene expression was confirmed by assessment of GFP fluorescence, and polyclonal GFP+ cultures were established by fluorescence activated cell sorting.

### Primary cell culture

Primary human leukaemia cells were sourced from the HTA licensed Barts Tissue Bank in compliance with the ethical approval as previously described^34^. Cells were assayed in a co-culture system using a GFP+ hTERT-immortalised human mesenchymal stem cell (MSC) feeder layer, given as a kind gift by Professor Dr D. Campana (St. Jude Children’s Hospital)^35^.

MSC cells at a maximum passage number of 10 were seeded into 96-well plates at a density of 8000 cells/well and allowed to adhere for 24 hours. Primary human bone marrow cells were then added directly onto the MSC layer without prior irradiation. Drug treatments were applied 24 hours after.

Following 48 h of drug exposure, non-adherent cells were removed and adherent cells were harvested using TrypLE Express dissociation reagent (Gibco). Cells were stained with Zombie Violet viability dye and a antibodies against AnnexinV-APC (BioLegend) and human-CD19-PE-Cy7 (BioLegend). MSCs were excluded from analysis using GFP expression. Leukaemic cell survival was quantified by normalising viable (Annexin V⁻/Zombie Violet⁻) human-CD19+ cells to vehicle-treated controls.

### Animal experiments

Experiments involving mice were performed with Queen Mary University of London veterinary oversight and complied with licensing requirements of the United Kingdom Home Office. C57BL/6 and NSG (NOD.Cg-Prkdc^scid^Il2rg^tm1Wjl^/SzJ) mice were purchased from Charles River Laboratories and housed in pathogen free conditions. Mixed female and male mice were used for patient derived xenograft experiments, while male mice were used for the p185^+^Arf^-/-^ dasatinib resistance model, based on previous engraftment and toxicity studies.

Patient derived xenografts were generated by tail vein injection of primary B-ALL cells. Individual mice were injected with 1-3 million cells and leukaemia development was monitored by intratibial bone marrow aspiration or tail vein phlebotomy. Engraftment was measured by flow cytometry and defined as mean tumour burden of at least 10% in bone marrow or 0.5% in blood. On confirmation of engraftment, animals were randomised to experimental arms and drugs administered in a blinded fashion. Pegargiminase was administered as a weekly intraperitoneal (i.p.) injection of 5 international units (equivalent to 575µg) diluted in 200ul warmed phosphate buffered saline, based on previous pharmacodynamic and pre-clinical studies^36,37^.

The p185^+^Arf^-/-^ mouse Ph+ ALL model was initiated and treated as previously described^29^. Each mouse was injected with 0.2 million viable cells on day 0, and bioluminescence imaging was performed twice weekly from day 4. Imaging was performed using an IVIS Lumina III (Revvity). Mice were shaved ventrally and injected i.p. with 100mg/kg Vivo Glo Luciferin (Promega) 10 minutes prior to acquisition, followed by anaesthesia with isofluorane. Dasatinib (10mg/kg) was prepared in 80mM citric acid as previously described^38^ and administered once daily by oral gavage from day 10, five days per week, with or without pegargiminase.

Mice were assessed for humane endpoints according to an objective scoring system, assessed daily by independent animal core staff.

### *ABL1* kinase domain sequencing

### *ABL1* kinase domain sequencing was performed as previously described^39^. Total RNA was extracted from mouse spleens and reverse transcribed to cDNA, which was then amplified using BCR::ABL1 e1a2 transcript specific primers. Library preparation was performed with the Illumina Nextera XT kit and paired-end sequencing was performed on an Illumina MiSeq platform. An in-house bioinformatics pipeline was utilised with a custom variant scoring tool for *ABL1* kinase domain variant identification.

### Western Blotting

Western blotting was performed as described previously^34^ using Cell Signalling Technology Cell Lysis Buffer to generate total cellular protein extracts. Representative blots from at least two independent experiments are shown.

### RT-qPCR

Real time quantitative PCR was performed as described previously^34^ using the QuantStudio 7 platform (Thermo Fisher). Analyses were performed using a minimum of 3 independent cDNA preparations from separate experiments, each performed in technical triplicate on a single plate. *GAPDH* was used for internal normalisation, except for *HBS1L* exon 1-2 inclusion ratio, which was calculated by normalising Ct values for primers spanning *HBS1L* exon 1-2 with the mean Ct values from primers spanning *HBS1L* exons 3-4 and 9-10. Primer sequences used are as follows: *HBS1L* exons 1-2 forward TCTACAGACTGGCCGTAGAGATCA, reverse CCCGGCATCGGAATGTT; *HBS1L* exons 3-4 forward TCGTCTTTATTCATGCCTTG, reverse TCCTTTTGCTATCTTTCCTG; *HBS1L* exons 9-10 forward TAAGCAAGCAGGTTTTAAGG, reverse TTTGTGAGTTCACTTGACTG; *ANPEP* forward CCTTCATTGTCAGTGAGTTC, reverse CAGCAAAGAAGTTAAGGATGG; *CD33* forward GCCTCATCTTCTTCATAGTG, reverse GTAACTTGGACTTCTTCTGG; *MS4A1* forward ATATACAACTGTGAACCAGC, reverse CCCAAGAACAGAGATTGTATG; *CD22* forward GGTTCTAGAATACTTGGCTTTC, reverse GATTGTGGAACAGGATGAAG; *GAPDH* forward CTCTCTGCTCCTCCTGTTC, reverse GGTGTCTGAGCGATGTGG.
