## Supplementary figures and images for "ASS1 deficiency defines a therapeutic vulnerability in Philadelphia chromosome-positive acute lymphoblastic leukaemia"

### supplemental_figures

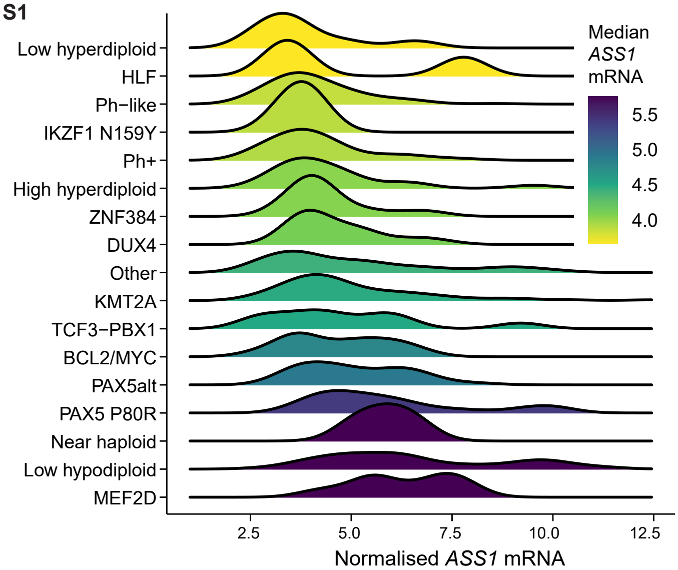

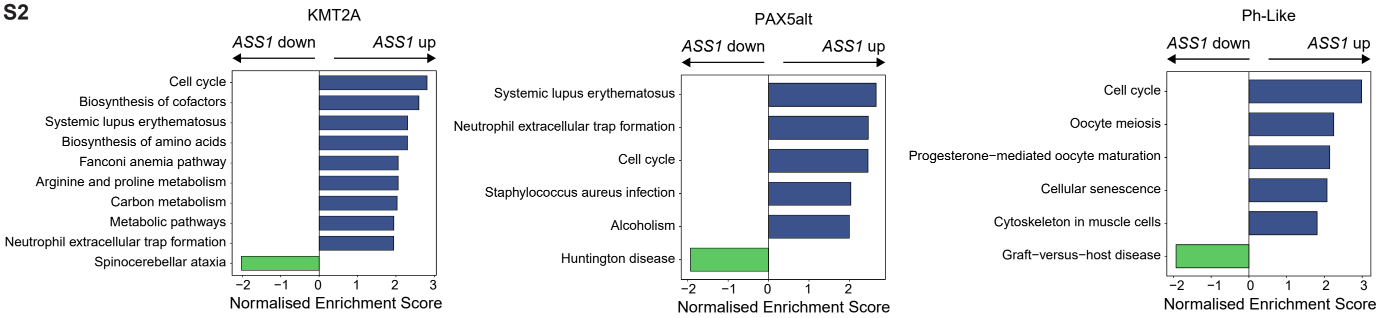

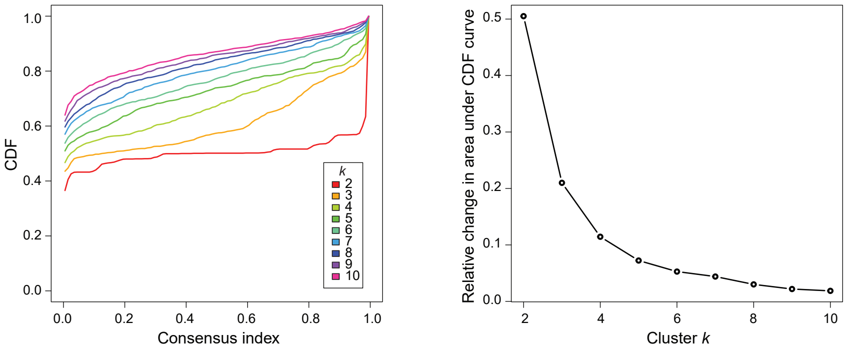

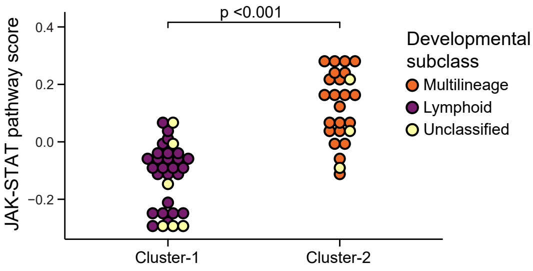

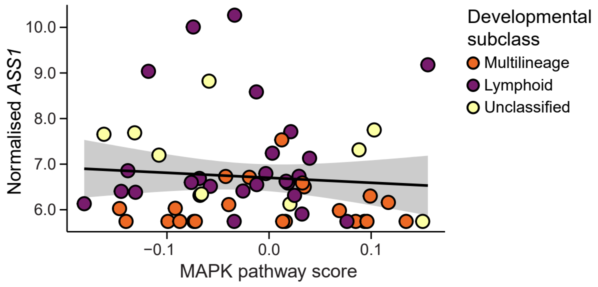

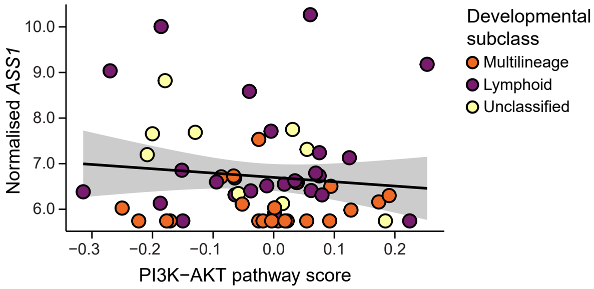

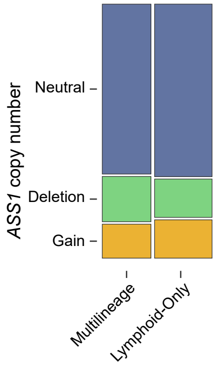

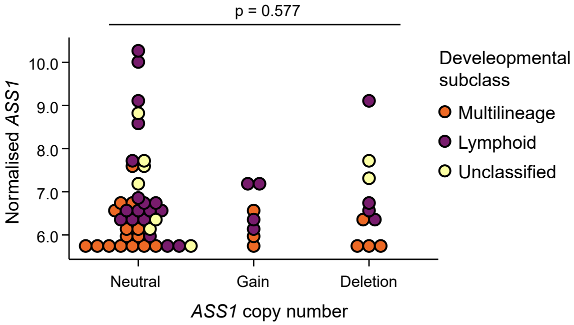

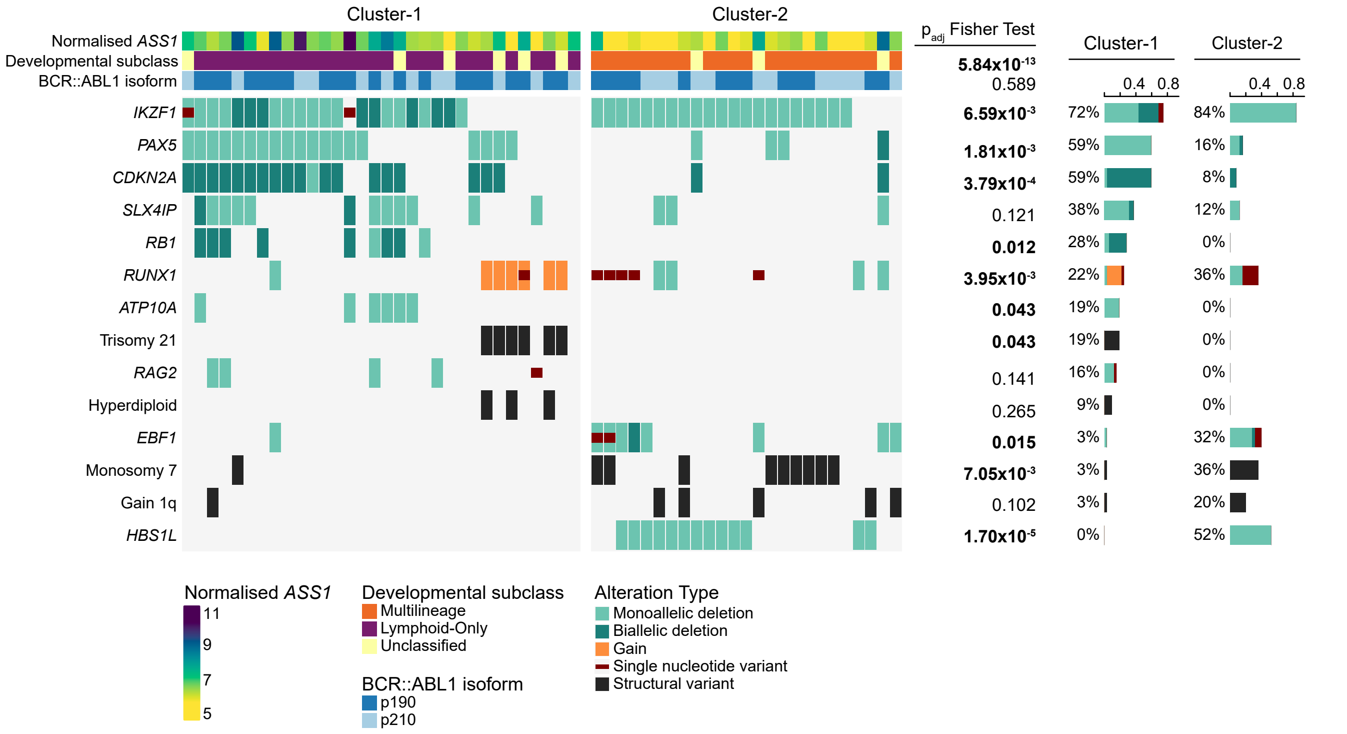

S10

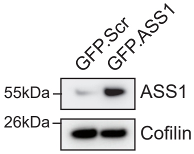


S11

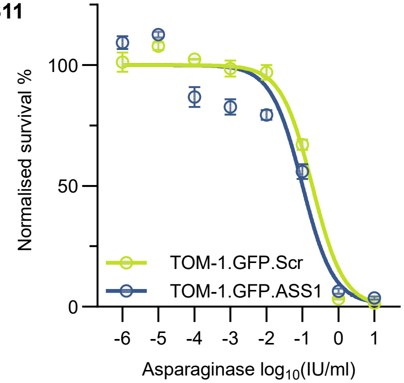


S12

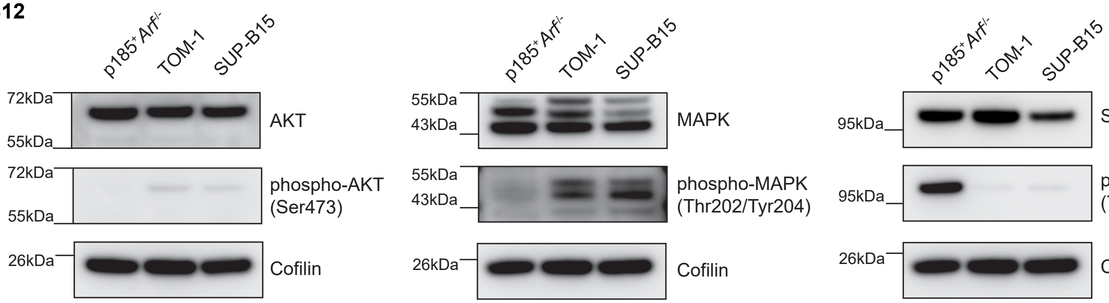


S13

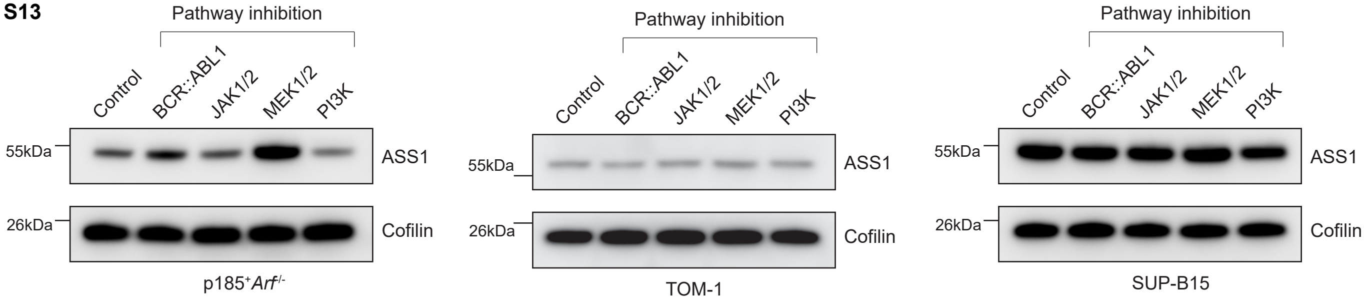


S14

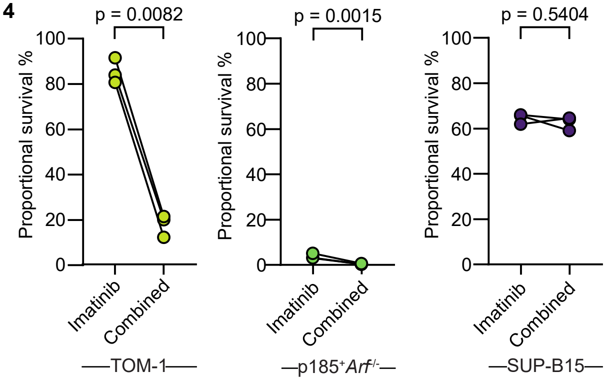

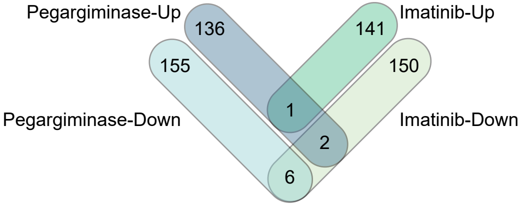

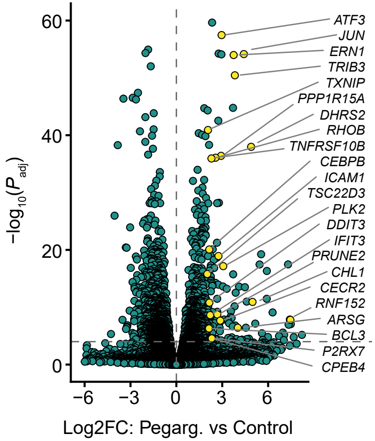
